# A Pathoconnectome of Early Neurodegeneration

**DOI:** 10.1101/2020.01.31.929265

**Authors:** Rebecca L. Pfeiffer, James R. Anderson, Jeebika Dahal, Jessica C. Garcia, Jia-Hui Yang, Crystal L. Sigulinsky, Kevin Rapp, Daniel P. Emrich, Carl B. Watt, Hope Morrison, Alexis R. Houser, Robert E. Marc, Bryan W. Jones

## Abstract

Connectomics has demonstrated that synaptic networks and their topologies are precise and directly correlate with physiology and behavior. The next extension of connectomics is pathoconnectomics: to map neural network synaptology and circuit topologies corrupted by neurological disease in order to identify robust targets for therapeutics. In this report, we characterize the first pathoconnectome, in this case, generated from a retina with photoreceptor degeneration. We observe aberrant connectivity in the rod-network pathway and novel synaptic connections deriving from neurite sprouting. These observations reveal principles of neuron responses to the loss of network components and can be extended to other neurodegenerative diseases.

## Role of connectomics in understanding neurological disease

The precise organization of neurons and their connections defines circuit topologies allowing normal nervous system function. This principal has been demonstrated in numerous neural networks from spinal circuits,^1^ to *C. Elegans*,^2^ to central nervous system (CNS),^3^ where distinct classes of neurons synapse with one another to create precise network specificities. Disruption of this organization leads to deficits or failures in neural network function.^4–6^ In a simple, static network, this is intuitive: Neuron A is presynaptic to neuron B, neuron B is presynaptic to neuron C, the loss of neuron A or B will lead to a lack of input to neuron C.^7^ Neural network complexity rapidly increases with the addition of more neurons and associated synapses. Understanding neural networks and how changes in them result in neurological disease is critical for understanding the progression of these diseases. Furthermore, creating targeted therapeutic interventions to stop, reverse, modify, or interact with degenerate neural networks will be greatly facilitated if we understand how networks are altered in disease.

In 2011, we published the first report of a connectome, (RC1).^8^ To date, more than 60 papers have expanded on the network motifs deriving from this retinal connectome.^8–11^ The retina offers a convenient substrate for studying mammalian neural networks and their complexities as it is compact and highly ordered, with a series of neuronal cell soma layers and plexiform or interconnecting layers containing repeating network motifs. Though annotating and analyzing the entire retinal connectome is not yet complete, we have a better understanding of the organization of its neural networks than we do of any other region of the mammalian CNS. The retina is also unique in its accessibility: gross histology and functional assays can be conducted non-invasively down to cellular resolution in living animals using optical coherence tomography (OCT)^12^ and electroretinogram (ERG)^13^ to track the progression of disease. In addition, recent work has demonstrated that retina behaves similarly to CNS with respect to late stage disease with many of the same features observed in retina as in other neuropathies of the brain.^14^

These features also make the retina an ideal candidate for mapping neuronal network disruptions that occur as a result of neurological disease by creating connectomes of pathological tissue: *Pathoconnectomes*. Here, we introduce the structure, features, and initial findings from Retinal Pathoconnectome 1 (RPC1) an open-access, serial-section transmission electron microscopy (ssTEM) connectome volume, generated from a rabbit model of autosomal-dominant cone-sparing retinitis pigmentosa (RP). Progressive neurodegenerative disease seen in retinal degenerative diseases like RP are initiated through a number of mechanisms. RP is a retinal disease that results in photoreceptor degeneration and blindness affecting approximately 1 in 4000 individuals. Although over 100 genes are associated with this disease, the sequence of degeneration in rod dominated RP is: rod photoreceptors degenerate causing neural retinal deafferentation, loss of night and peripheral vision, followed secondarily by the degeneration of cone photoreceptors and total or near total vision loss.^15^ In the retina, these negative plastic changes and cell death revise the circuit topology through a process called retinal remodeling. This report examines the changes in circuitry of the retina resulting in *rewiring*, which is the specific process of changes in network topology that occur through neurite outgrowth, formation of new synapses, and loss of existing connections. For further review of photoreceptor degenerations and retinal remodeling, see Jones and Marc, 2005.^16^

## Volume assembly and small molecule tagging

Ultrastructural connectomes provide the high-resolution capabilities of transmission electron microscopy (TEM) with the quantitative aspects of computational molecular phenotyping (CMP) to molecularly label cell identity.^8^ These approaches have been previously used to create a healthy rabbit connectome RC1: a 250μm in diameter volume of retina spanning from the apical inner nuclear layer (INL) to the basal ganglion cell layer (GCL), which serves as a ground truth for normal retinal circuit topology. RPC1 was constructed in identical fashion. In electron microscopy, adequate resolution is required to definitively identify all network components (2.18nm/px is the lowest resolution in which gap junctions can be positively identified) and the identification of pre- and post-synaptic structures is crucial to the accurate identification of networks. Therefore, our routine TEM acquisitions are acquired at 2.18 nm/pixel, and sections can be re-imaged at 0.27 nm/pixel, with a goniometric tilt as necessary to confirm certain circuit identities like gap junctions^9,10,17^. Annotation of every chemical or electrical synapse type is made to avoid false positives of connectivity and minimize missed structures. For further description of synapse identification metrics used in this study, see supplementary figure 1.

Along with ultrastructural data, molecular tagging of cellular identity is fundamental to understanding contributors to neuronal networks. CMP allows for cell classification through quantitation of small molecules comprising metabolic components.^18,19^ Small molecule IgGs to GABA, L-glutamate, L-glutamine, glycine, and taurine, in addition to IgGs targeting glial fibrillary acidic protein (GFAP) were used in the classification of cells in the RPC1 volume. Following volume assembly, navigation and annotation of the dataset is performed in the Viking software environment.^8^ From a Viking-annotated dataset, we can render 2D and 3D morphologies of cells including their subcellular components (e.g., pre- and postsynaptic densities, organelles, gap junctions, etc.), as well as use custom (VikingView and cell sketches) and commercial software (Tulip and Blender) packages to evaluate and visualize networks.^9^

## Retinal pathoconnectome 1 (RPC1)

RPC1 is composed of 946 serial TEM sections with 14 intercalated CMP probes (Figure 1 (larger versions of all figures are also provided at the end of the manuscript)). Because 1 out of 30 sections cut were reserved on slides, we have 17 unstained sections available for future probes. The volume itself is narrower in diameter (70μm) than RC1 (250μm), allowing for more rapid capture and assembly, while large enough to capture repeated rod-network motifs. RPC1 is the first pathoconnectome volume in a series of 3, each with increasingly progressive photoreceptor degeneration/neurodegeneration to chronicle landmarks in rewiring as the disease progresses. The first pathoconnectome, RPC1 derives from a 10-month-old transgenic (Tg) P347L rabbit retina, model of RP.^20,21^ In the selected region of retina (peri-visual streak), photoreceptor degeneration has initiated and the outer nuclear layer (ONL) is reduced in thickness by approximately 50%. However, though rod photoreceptors are the most heavily degenerated, many are still intact. The remaining rod photoreceptors, along with the entire bipolar cell population, allow for investigation of early-stage RP rewiring. In total, 321 neuronal and glial somas have been identified within the volume and assigned to broad classes in RPC1. The breakdown of cells is as follows: 113 photoreceptors, 5 horizontal cells, 81 amacrine cells, 79 bipolar cells, 4 ganglion cells, at least 36 Müller glia, and 4 microglia. Examination of these cell classes has revealed altered network topologies associated with rewiring in early photoreceptor degeneration, *prior to* complete rod degeneration or inner retinal neuronal loss.

**Figure 1:**
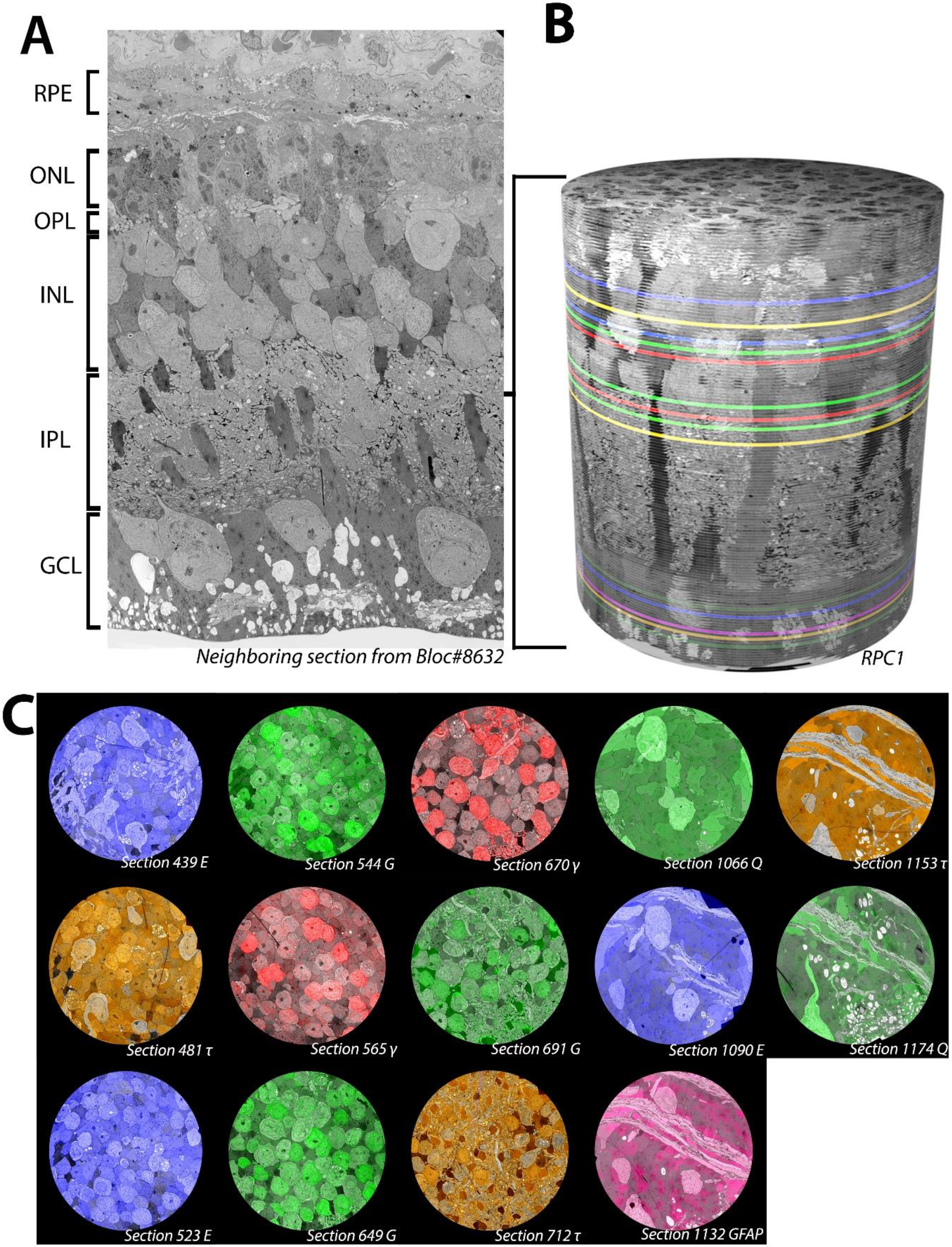
Overview of RPC1. (A) Vertical section through tissue directly adjacent to the tissue processed for the RPC1 volume. (B) 3D composite volume of RPC1. Pseudocolored sections illustrate the locations of CMP sections for small molecules indicated in 1C. (C)Overlaid CMP sections from RPC1 on their adjacent TEM sections.

## Rod-network pathologies

The first network we sought to characterize in RPC1 was the rod-network, due to the rod degenerative phenotype of our model system. The known literature of complete retinal networks is vast, but a simplified graphical description of the rod network and its associated components can be summarized here (Figure 2). For more comprehensive descriptions of additional retinal networks see: Marc et al., 2014, Marc and Sigulinsky et al., 2018, Diamond, 2017, Thoreson and Dacey, 2019, and Lauritzen et al., 2019 ^9,10,22,23^

**Figure 2:**
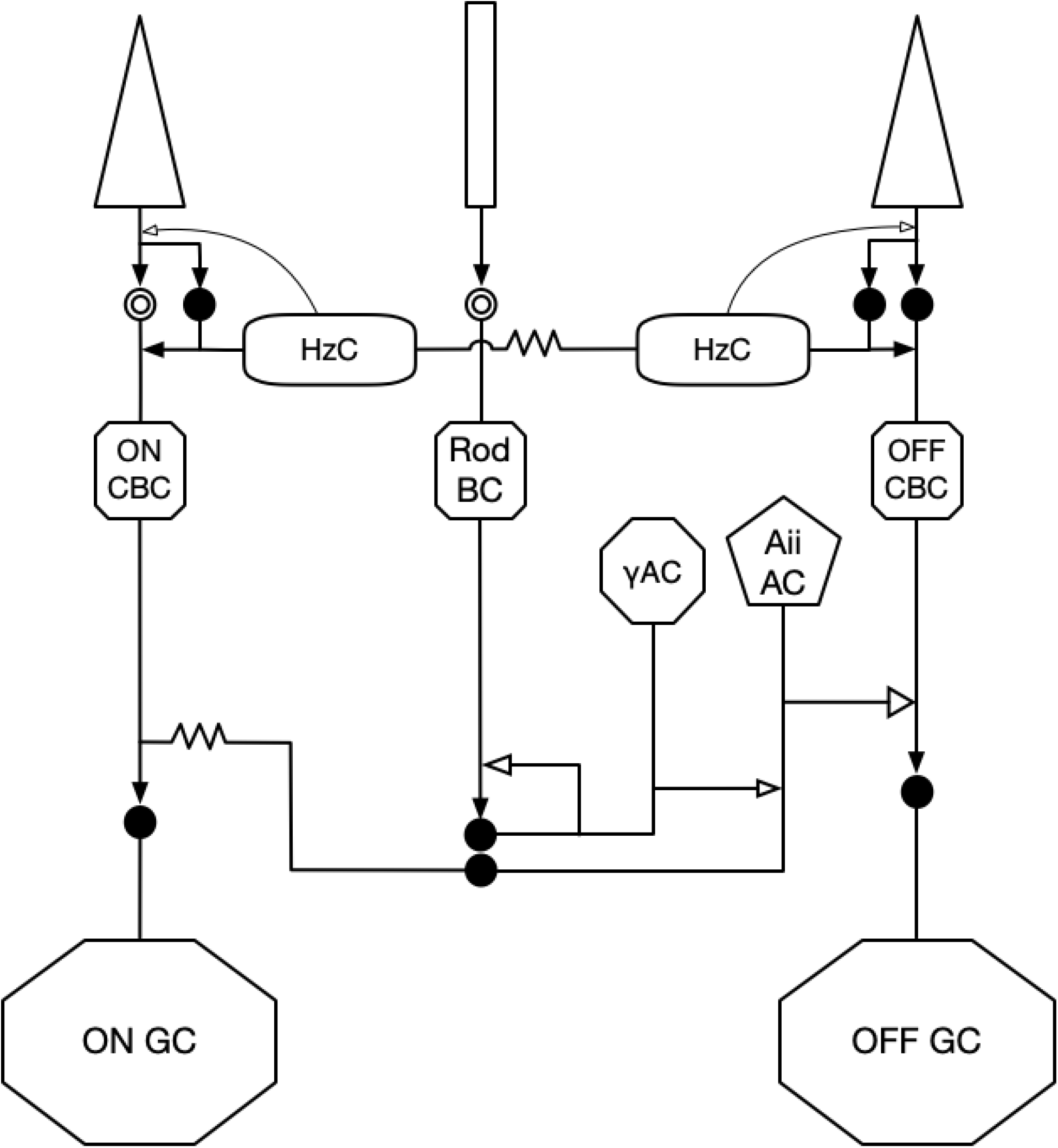
A simplified diagram of RodBC network. Black arrows indicate glutamatergic synapses. Open arrows indicate inhibitory synapses (GABA or glycine). Solid circles indicate ionotropic glutamate receptors, while mGluR6 is indicated by the double open circles. Gap junctions are indicated by zig-zag lines

Rod photoreceptors (rods) are specialized neurons responsible for detection of photons in low-light (scotopic) conditions. Following photon detection, rods initiate an enzymatic cascade ultimately leading to a depolarization, and then synaptically signaling to postsynaptic rod bipolar cells (RodBCs), via the first synapse in vision located in the outer plexiform layer (OPL). Axons of RodBCs branch and synapse in the deepest layer of the inner plexiform layer (IPL), close to, but not connecting with ganglion cells in the ganglion cell layer (GCL). RodBCs are presynaptic to at least 2 classes of amacrine cells: GABAergic amacrine cells (γACs) and Aii glycinergic amacrine cells (Aii GACs). Aii GACs are chemically presynaptic to OFF cone bipolar cells (OFF CBC) using the neurotransmitter glycine, causing inhibition. However, in the ON-layer, Aii GACs make gap junctions with ON cone bipolar cells (ON CBCs), sharing their polarization state through electrical connections with the ON CBCs. ON CBCs in turn pass rod signals to ganglion cells that ultimately project out of the retina, and into the brain via the optic nerve. In healthy retina, RodBCs are never presynaptic to ganglion cells and therefore “piggyback” on the ON CBC pathway using the Aii GAC as described above. This rod pathway is highly conserved and observed in all mammalian retinas.

RPC1 contains 17 RodBCs, 8 of which have axonal and dendritic arbors contained entirely within the volume. RodBCs are traditionally thought to be exclusively postsynaptic to rod photoreceptors in the healthy retina. Recent studies have proposed that RodBCs may be post-synaptic to cones in addition to rods in healthy retina, though the resolution and lack of clear post-synaptic densities to the cone pedicles make it difficult to determine whether the RodBCs are truly synapsing with the cone pedicle or merely traversing in close proximity^24,25^. Because our RC1 volume does not include an OPL, we cannot directly assess whether these contacts exist using our identification metrics, though the serial TEM reconstruction of RodBCs in a healthy mouse by Tsukamoto and Omi failed to identify any cone inputs to these cells.^11^ In photoreceptor degeneration retinal diseases however, it is widely accepted that RodBCs retract their rod-contacting dendrites as rods die and extend new neurites towards cone pedicles making peripheral contacts.^26^ In evaluating the OPL of RPC1, we find this process initiates prior to complete rod degeneration (Figure 3). In early retinal degeneration, RodBCs maintain contact with existing rods, while simultaneously making aberrant contacts with cones. We observed 10 RodBCs post-synaptic to cone photoreceptors, with 4 of those conecontacting RodBCs maintaining at least 1 post-synaptic density to a rod axon terminal (spherule). Though most cone-contacts are with traditional cone axon terminals (pedicles), we find one instance of a sprout off the primary pedicle with a secondary terminal presynaptic to RodBC 822. We find an additional 7 RodBCs with densities opposing non-traditional structures containing ribbons. In these cases, we find 1 RodBC opposing axonal ribbons and no terminal, and 6 opposing ribbon terminals inconsistent with morphology typically associated with rod spherules or cone pedicles and cannot be conclusively identified. Because 9 of the RodBC dendritic arbors are not entirely contained within the volume, this indicates the number of cone contacts made by RodBCs in RPC1 may be greater than the currently tabulated 58.8% (10/17). These results confirm previous light microscopy work indicating RodBCs extend neurites to cone pedicles as the rod photoreceptors degenerate, and expands upon this by determining RodBCs extend neurites towards cone pedicles prior to becoming fully deafferented from rod spherules. Although, a prior study demonstrated cone neurite sprouting in late retinal degeneration following substantial cone degeneration,^27^ we find this process initiates early in photoreceptor degeneration, prior to the complete loss of rod photoreceptors and onset of cone degeneration.

**Figure 3:**
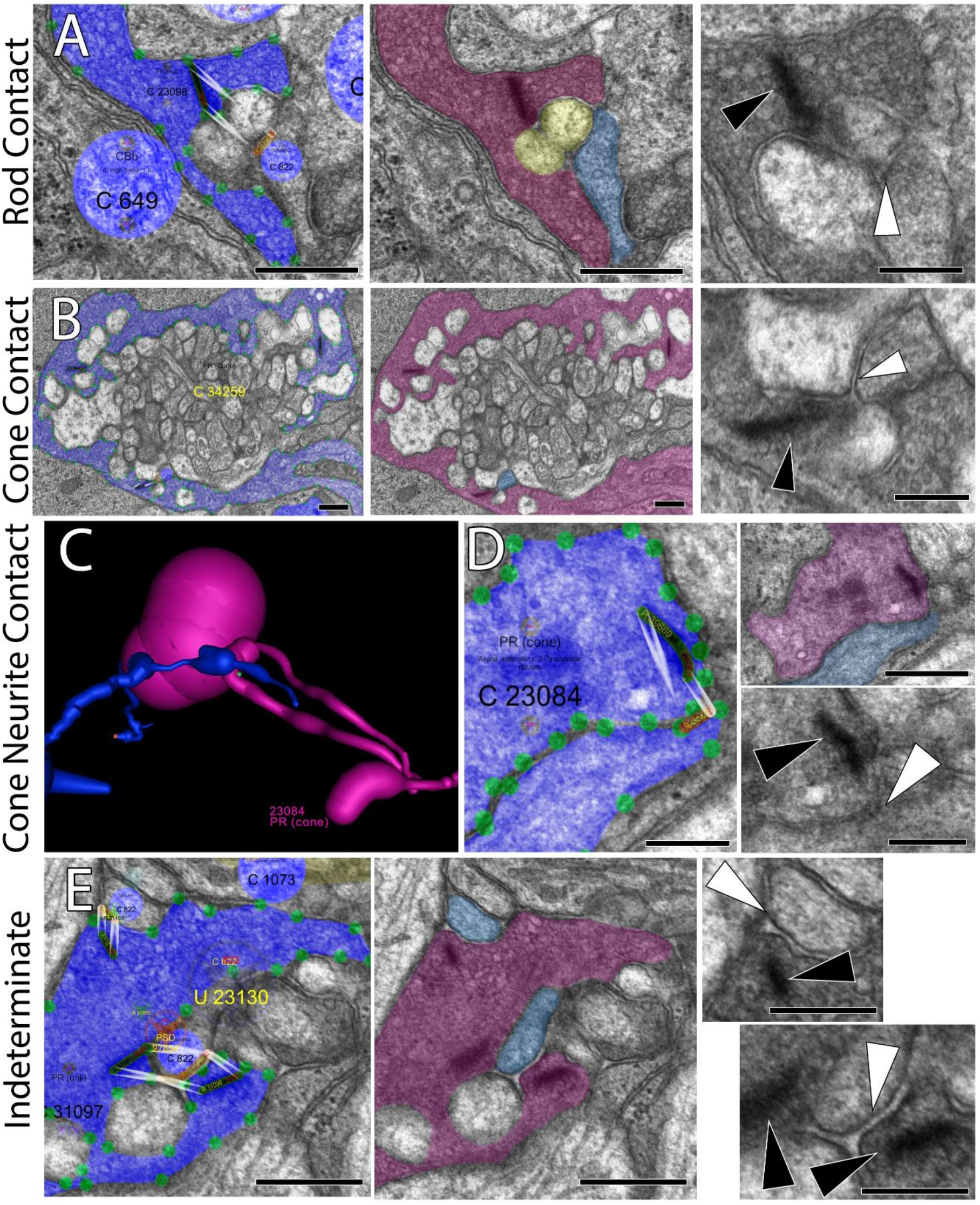
RodBC dendritic contacts. (A) Representative rod photoreceptor synapse of RodBC 822 postsynaptic to rod. (B) Representative cone photoreceptor synapse between a cone photoreceptor and RodBC 1223. (C) 3D rendering of cone photoreceptor 23084 main terminal and 2 neurite processes. (D) Representative photoreceptor synapse between a cone neurite ribbon and RodBC 822 (E) Representative photoreceptor synapses between an indeterminate photoreceptor and RodBC 822. Black arrowheads indicate ribbon densities, while white arrowheads indicate associated post-synaptic densities. Viking Annotation and Pseudocolored Images Scale bars 500nm. Higher Magnification Scale Bars 250nm

We then compared the inner retinal network of RPC1 with the previously described network topologies of RC1. The total ribbon synapses made by complete RodBCs of RPC1 (n=8) are similar to representative RodBCs (n=5) from RC1. RPC1 ribbons: 34.125 ± 4.29 (mean ± standard deviation) and RC1 ribbons: 34.4 ± 2.5 (p=0.8, unpaired Student’s-t-test). The prevalence of postsynaptic densities was also similar between RPC1: 69.625 ± 14.050 and RC1 62.8 ± 13.498 (p=0.406, unpaired Student’s-t-test). These results suggest that chemical synapse numbers are not changed in number at this stage of photoreceptor degeneration.

Chemical synapses between RodBCs, Aii GACs, and yACs appear to be intact in the early degenerate retina and occur in similar frequency to that observed in healthy retina. However, we observed the emergence of gap junctions involving RodBCs of RPC1. In the adult healthy mammalian retina, RodBCs do not make gap junctions. This is confirmed in our RC1 volume, where none of the 104 RodBC axonal arbors make a single gap junction (Sigulinsky et al., submitted). In contrast, all 17 RodBCs found within RPC1 form at least 1 gap junction. In total, we have identified 50 candidate gap junctions, 18 of which were confirmed using high magnification (0.43nm/pixel) and goniometric tilt (Figure 4). At least one gap junction per RodBC was confirmed using high magnification with the exception of RodBC 26167, which is largely off volume and recapture was not possible of its only identified candidate gap junction. Next, we identified coupled partners. No gap junctions are found between paired RodBCs in RPC1, despite substantial overlap of arbors in some areas (Figure 4C). Of the 50 gap junctions, 48 (96%) are positively identified with Aii GACs. In all cases of gap junctions with Aii GACs in RPC1, the individual RodBC is also heavily presynaptic to the same Aii GAC via glutamatergic ribbon synapses, often within the same varicosity as the gap junction. Of the gap junctions not made with a confirmed Aii GAC, 1 is with an unidentified partner due to being near the volume edge and the other is with an ON CBC. These data demonstrate that RodBCs undergo rewiring with respect to their photoreceptor input prior to complete rod degeneration and simultaneously alter their inner retina synaptology through the formation of gap junctions, especially with Aii GACs. There also appears to be preferred class-specific partnering via gap junctions that may reveal the presence of cell-cell markers or targets for therapeutic intervention. What implications this has for network performance or stability is currently unclear, but is an area of future modeling work.

**Figure 4:**
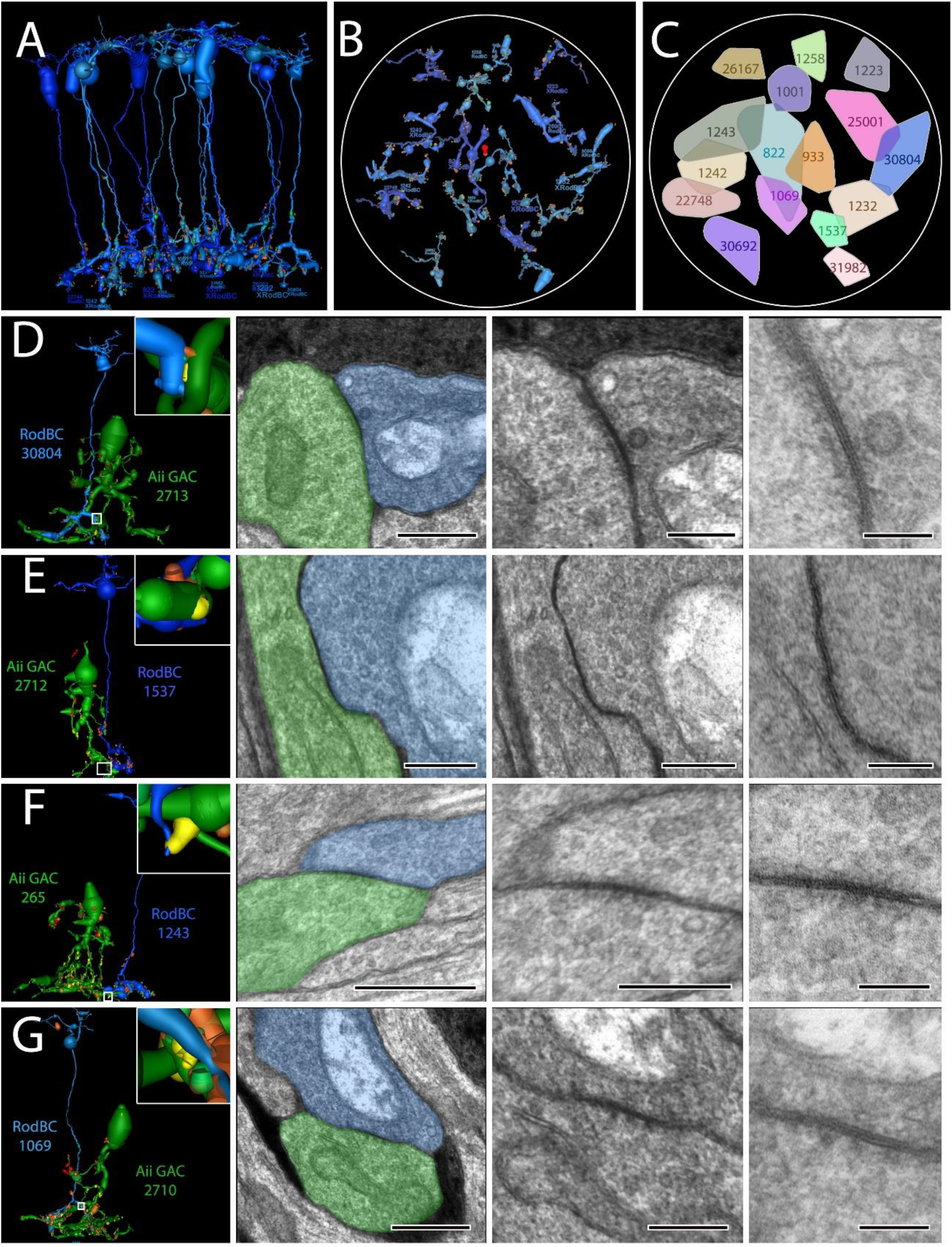
RodBC gap junctions with Aii GACs (A) 3D rendering of all 16 RodBCs from RPC1. (B) RodBC axonal arbor fields in RPC1. (C) Convex Hulls of all RodBCs in RPC1 (D-E) Gap junctions between RodBCs and Aii GACs. Left panel is 3D rendering with inset higher magnification image of specific gap junction annotations. White boxed image corresponds to white box in adjacent left panel. * indicate ribbons and # indicates gap junctions arising in same varicosity. 100nm scale images are 25k recaptures of gap junction structures

## GABAergic amacrine cell neurite sprouting

RPC1 has 31 identified GABAergic amacrine cell (γACs) somas contained within the volume, that were also annotated. In the healthy retina, γACs have a soma that is predominately positioned just apical to the IPL, with processes extending solely into the IPL, and is consistent with findings in the normal retinal connectome, RC1 which has 107 GABAergic amacrine cell somas. In disease, prior work has demonstrated that processes of γACs can be found extending apically towards the OPL during photoreceptor degeneration.^20^ However, it was unclear whether these GABAergic processes were neurites that extended off of the normal processes below the soma, or whether they arose from the soma itself. In RPC1, 4 of the 31 γACs extend processes from the apical side of their somas into the OPL (γAC cell #s: 769, 993, 997, and 2627) (Figure 5). Not only is this confirmatory at ultrastructure of the previously described novel GABAergic processes in the OPL, but demonstrates that the extension of these processes arises from the soma, and manifests prior to complete loss of rods. Connectomics has the additional advantage of permitting identification of the synaptic partners of these aberrant processes. The summary of these aberrant processes connectivities is as follows: γAC 769 extends into the OPL, but makes no identifiable synapses. γAC 993 is the most complex of these apical extending processes making 6 synaptic contacts in the OPL: 2 postsynaptic densities to conventional synapses originating from horizontal cells and 4 synapses in which γAC 993 is presynaptic to multiple classes of ON bipolar cells. γAC 997 is postsynaptic to a single horizontal cell, and γAC 2627 is presynaptic to a horizontal cell and an OFF bipolar cell. These connectivites are novel in 2 distinct ways: 1) γACs in RPC1 make typologically correct contacts with bipolar cells, even when encountered in the wrong lamina. 2) γACs make aberrant synapses with horizontal cells that they never come in physical proximity with in the healthy retina. This demonstrates the ability of deviant processes that infiltrate an aberrant layer to make synapses with numerous typologically correct and incorrect partners, clearly altering the neural networks downstream of cells actively undergoing degeneration.

**Figure 5:**
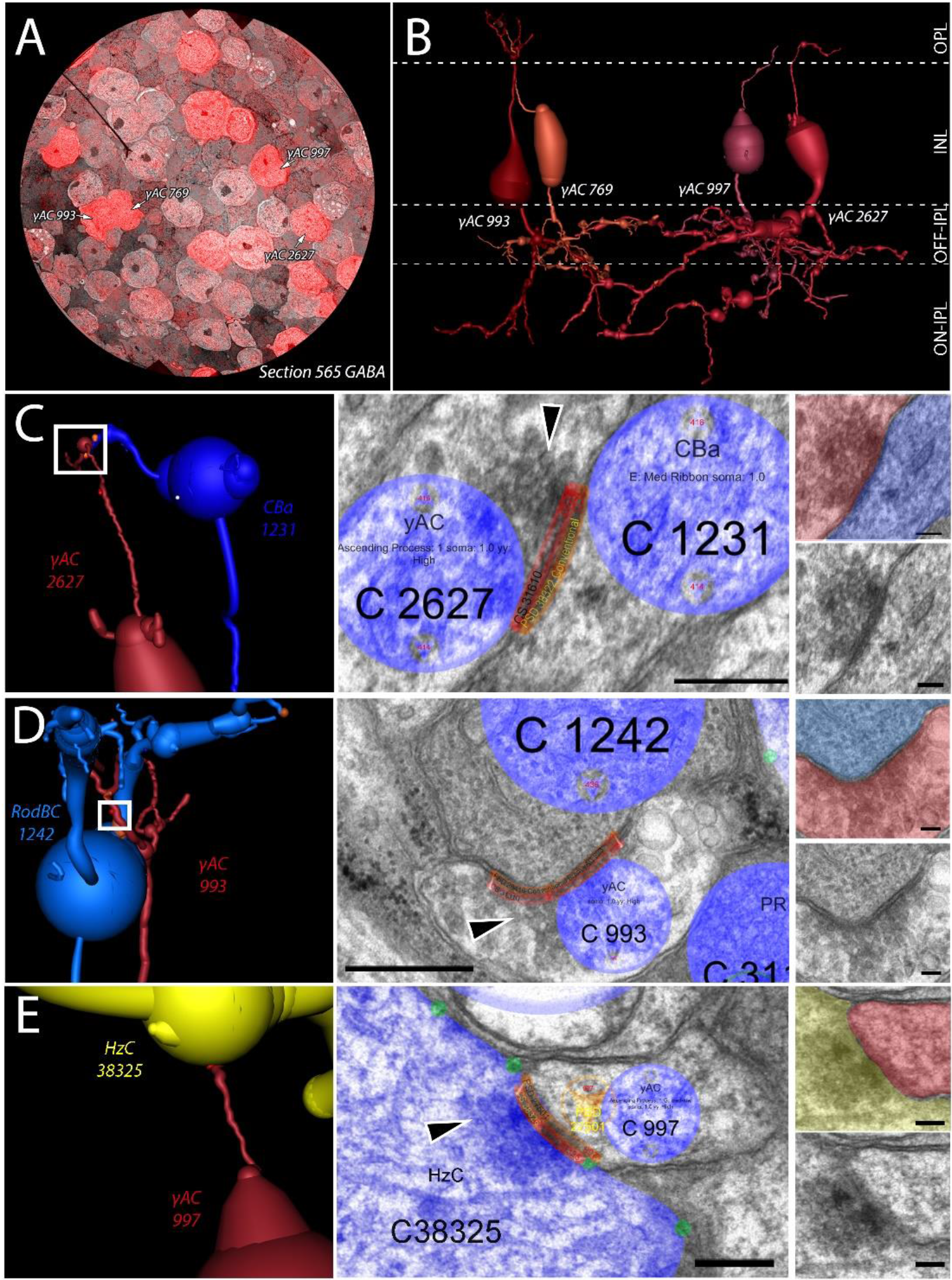
Ascending process of GABAergic amacrine cells. (A) CMP overlay of GABA section 565 on its neighboring TEM section. Intensity of red color indicates higher levels of GABA contained within the cell. (B) 3D rendering of the 4 γACs with ascending processes in RPC1. (C) Synapse between γAC 2627 and OFF-BC 1231 in the OPL. White box indicates region of synapse shown on the right in increasing magnification. Black arrowhead indicates the presynaptic vesicle cloud. (D) Synapse between γAC 993 and RodBC 1242 in the OPL. White box indicates region of synapse shown on the right in increasing magnification. Black arrowhead indicates the presynaptic vesicle cloud. (E) Synapse between γAC 997 and HzC 38325 in the OPL. White box indicates region of synapse shown on the right in increasing magnification. Black arrowhead indicates the presynaptic vesicle cloud. Scale bars: 250nm (Viking annotated) 100nm (smaller inset).

## Neural network rewiring will affect therapeutics

Rewiring of neural networks through plasticity is a prominent consequence of injury or disease in the central nervous system. Multiple studies have explored the mechanisms of neurite outgrowth,^28^ localized the sites of projections,^26,29^ and identified a subset of synaptic partners.^29–31^ Unfortunately, exploring this using antibody probes relies on pre-existing expectations of proteins that may be involved. In addition, even when the correct protein is predicted, many synaptic structures underlying networks are too small to resolve in light microscopy or lower resolution electron microscopy approaches. Ultrastructural pathoconnectomes captured at 2nm/pixel allows for exploration and characterization of these novel networks formed by native neurons and aberrant neurites to establish precise wiring topologies, as well as identify potential rulesets that neural tissues undergoing degeneration may adopt. These networks also identify targets for potential therapeutic intervention.

In this example of early progressive neurodegeneration, this region of retina would likely still exhibit both day and night vision, but might have difficulty adapting between the two environments given corruption of the Aii related network. These findings support clinical observations in patients undergoing early stages of progressive retinal degenerative diseases. Our observations of changes in the rod pathway and γAC connectivities are important contributing components to this deficit, but are occurring prior to substantial inner retina neuronal cell loss and certainly prior to the network completely failing. RodBCs extend processes away from rod photoreceptor terminals (while maintaining contact with others) and contact cone pedicles in addition to apparent neurites off of cone pedicles. Apical neurites from γACs, not only extend and traverse the OPL, but also make anatomically confirmed synapses with bipolar cells in addition to horizontal cells, which they never encounter in healthy retina. These results identify the neurons involved in early neurite outgrowth and what cells are postsynaptic to these aberrant processes, leading to further questions about cell to cell recognition in plasticity and how it can might be controlled experimentally and therapeutically.

Specificity in synaptic partners is critical to the function of a neural network. Many complex feedforward and feedback loops have emerged as underlying properties long observed in physiological studies. For example, psychophysical studies of vision prompted many hypotheses of crossover inhibition between networks, which were later specifically defined using connectomics^9^. The present pathoconnectome reveals the early onset of many network-level changes involved in the breakdown of primary visual pathways within the retina (Figure 6). The emergence of gap junctional coupling between RodBCs and Aii GACs demonstrates the deterioration of the rod-network prior to the complete degeneration of the rod photoreceptors and likely compromises the spatial and temporal resolution found in the intact network.

**Figure 6:**
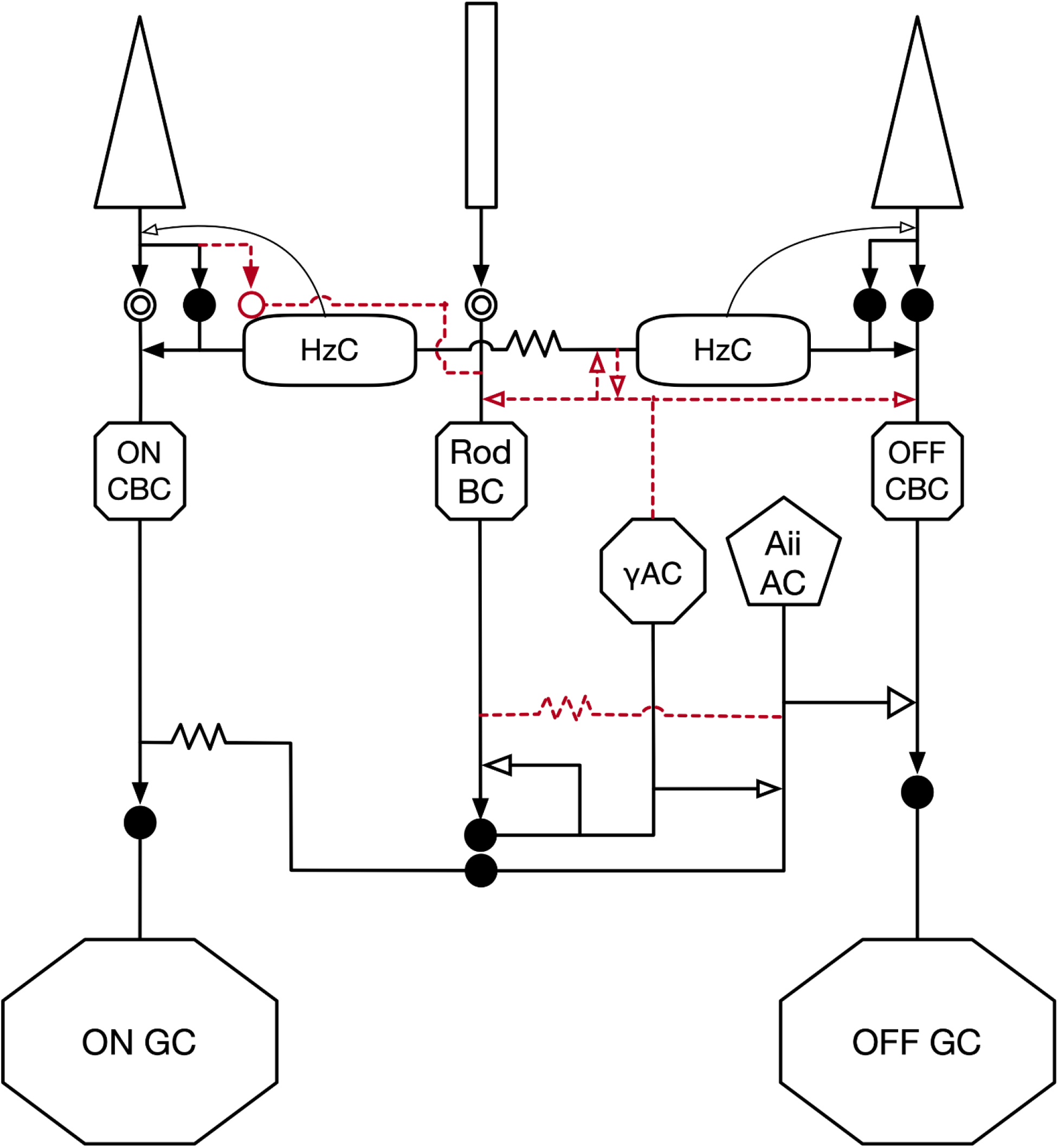
Altered network found in RPC1. Black arrows indicate glutamatergic synapses. Open arrows indicate inhibitory synapses (GABA or glycine). Solid circles indicate ionotropic glutamate receptors, while mGluR6 is indicated by the double open circles. Gap junctions are indicated by zig-zag lines. Red lines indicate aberrant connectivities observed in the RPC1 volume.

## Pathoconnectomes provide a map for therapeutics

This study introduces the first ultrastructural pathoconnectome and demonstrates applicability to understanding rewiring in neurological disease. Furthermore, it demonstrates the importance of using connectomics approaches to describe pathological consequences for neural network disruption through injury or disease. As future pathoconnectomes explore more advanced stages of neurodegeneration, we will learn more about the progression, what ultimately causes networks to fail, and describe potential therapeutic intervention targets. These principles are likely to be directly applicable to the broader CNS, as retinal remodeling exhibits the same components as CNS negative plasticity in disease, as well as similar proteomic findings observed in neurodegenerative diseases like Alzheimer’s, Parkinson’s and others.^14^

## Acknowledgements

This work was supported by the National Institutes of Health [RO1 EY015128(BWJ), RO1 EY028927(BWJ), P30 EY014800(Core)]; and an Unrestricted Research Grant from Research to Prevent Blindness, New York, NY to the Department of Ophthalmology & Visual Sciences, University of Utah

## Author Contributions

RL Pfeiffer, BW Jones, and RE Marc designed the experiment, selected the tissue, and oversaw the generation of the RPC1 volume and annotations. RL Pfeiffer and BW Jones wrote the manuscript, CL Sigulinski provided extensive edits. RL Pfeiffer and JR Anderson assembled the volume from initial TEM acquisitions. JR Anderson designed, wrote, and supports the Viking software environment. RL Pfeiffer, J Dahal, JC Garcia, JH Yang, CL Sigulinski, DP Emrich, H Morrison, AR Houser, RE Marc, and BW Jones annotated the dataset. K Rapp did the TEM imaging and high magnification recaptures. CB Watt stained the dataset. JH Yang manually sectioned the tissue for generation of the volume. RL Pfeiffer preformed all of the CMP for incorporation into the volume.

## Methods

### Volume Assembly

Tissues are fixed in a mixed aldehyde solution (1% formaldehyde, 2.5% glutaraldehyde), dehydrated in graded alcohols, then embedded in epoxy resins. Tissues are then serially sectioned at 70nm and prepared for TEM capture or CMP analysis by placing the section on a formvar grid or 12-well slide, respectively. Serial TEM images were acquired on a JEOL JEM-1400 electron microscope with a 16-Mpixel Gatan camera, while CMP images were acquired using a Leica light-microscope affixed with 8-bit CCD camera. RPC1 was assembled using the custom Nornir tools^1^ developed for volume assembly. Nornir combines individual images acquired from transmission electron microscopy or from a light-microscope, and registers individual image tiles into assembled mosaics before using automated registration to align adjacent sections throughout the volume, with minimal manual correction.^1,2^

### Computational Molecular Phenotyping (CMP)

CMP quantitatively evaluates the combinations of small molecules, which are stoichiometrically trapped during fixation,^3–5^ in addition to glutaraldehyde/osmium-tolerant IgGs. The concentrations of the different amino acids and proteins are used in the identification of cells beyond their morphology and synaptology (Table 1). For more in depth information on CMP technologies and usage see Marc, et al., 1995.^4^

**Table 1:**
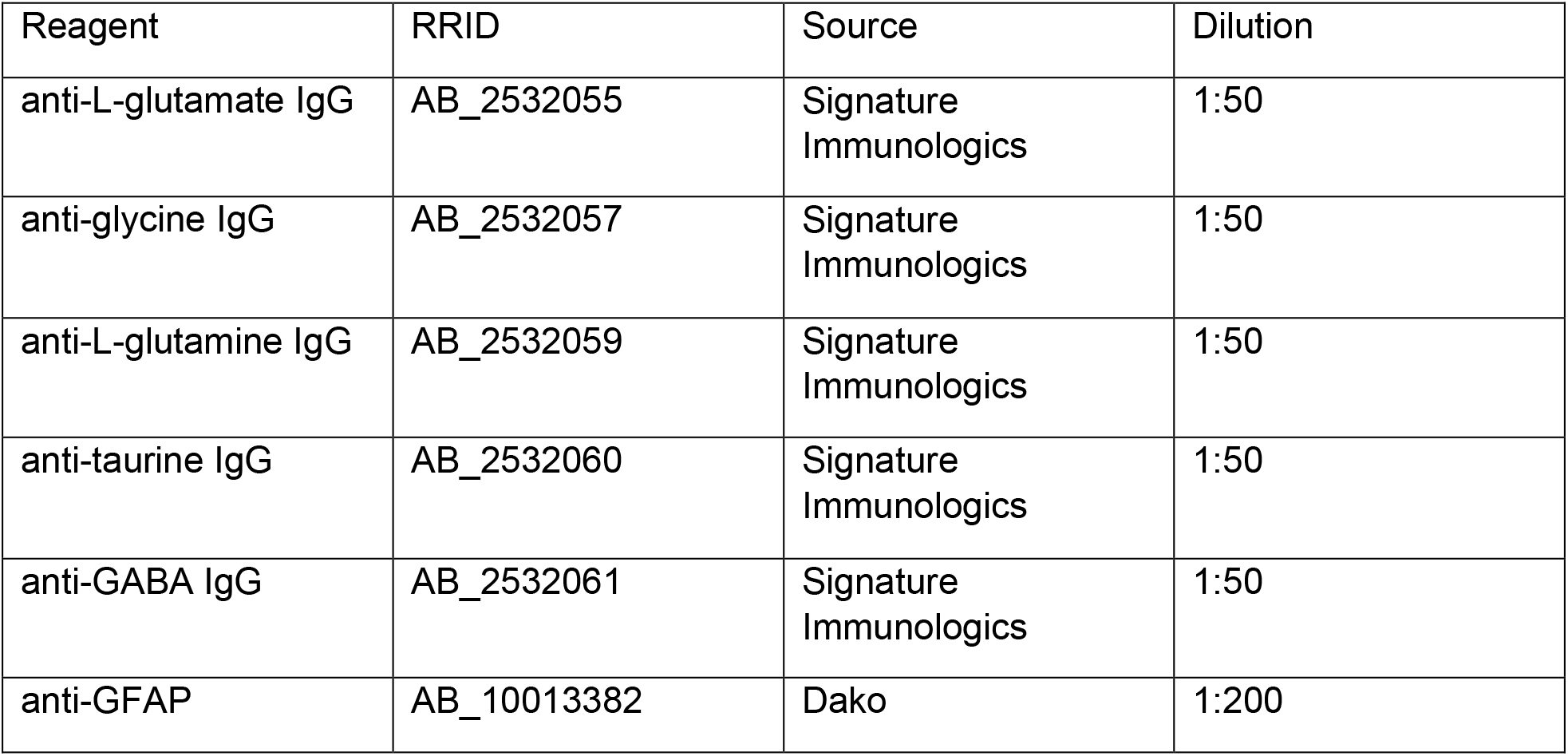

### Viking Software

Viking annotations encode information about the size, structure, and location of all annotated structures facilitating database-base queries of parameters including statistics of encoded structures.^2,6^ These databases are publicly accessible at www.connectomes.utah.edu and http://websvc1.connectomes.utah.edu/RPC1/OData.

### Figure Assembly

Figures were assembled using screenshots from the RPC1 volume in Viking (http://connectomes.utah.edu/, RRID:SCR_005986), 3D renderings of cells were generated in VikingView (DOI: 10.5281/zenodo.3267451, https://zenodo.org/record/3267451#.XSUW1OhKguU), and whole volume renderings were generated in Blender (http://www.blender.org/; RRID: SCR_008606). Final figure assembly was done in Photoshop 6.0.

## Footnotes

1 https://github.com/jamesra/nornir-buildmanager

