## Supplemental Information for "A Pathoconnectome of Early Neurodegeneration"

One of the primary challenges of connectomics is the correct identification of connections between cells in a volume of neural tissue. Many modalities can be used in this effort, and each have specific advantages and limitations, which we will broadly outline with a more comprehensive recent review of each technique cited. On a macro-scale functional magnetic resonance imaging (fMRI) and diffusion tensor imaging (DTI) are used with the primary advantage of being applicable to the entire brain of living animals, including humans. Its primary disadvantage is the lack of resolution: only large axon tracts or regions of activation can be observed and mapped.[1] On a meso-scale, tract tracing is employed. It is also amenable to tracing neuronal processes over long distances, with the added advantage of targeting of specific networks using a variety of in vivo and histochemistry techniques.[2] The primary disadvantage of tract tracing is the need for specific injections, dyes, or antibodies to label the individual tracts, and it also suffers from problems with resolution in especially dense networks. In the micro-scale of connectomics various modalities of serial section electron microscopy are employed, most commonly serial block face scanning electron microscopy (SBFSEM), and serial section transmission electron microscopy (ssTEM).[3] ssTEM has the primary advantage over techniques of high resolution with relatively short capture times. We routinely capture at 2.18nm/px and archive grids so that specific structures of interest can be reimaged at up to 0.25nm/px. Though we find ssTEM the best technique for complete, accurate, ultrastructural connectomics, ssTEM also has its limitations. One limitation is size, in that tissues of interest have to fit on standard formvar coated grids. The other primary difficulty is the fragility of grids through handling, where they can break occasionally even with expert handling in sectioning, staining, and imaging. SBFSEM has the advantage of not requiring sections to be cut and imaged individually, this leads to less need for section-to-section alignment and fewer specialized personnel.[4] The disadvantages of this SBFSEM are that the sections are discarded as they are cut, so recapture of any sections is impossible. The other major disadvantage of SBFSEM is resolution for large scale captures. Routine SBFSEM is done at a resolution of 10-30nm/px. At this resolution, detection of synapses can only be accomplished by finding large pre-synaptic vesicle pools and post synaptic membranes with higher density than their surrounds and inferring the synapse. This resolution fails to resolve many smaller synapses and all gap junctions in their entirety. What cannot be seen cannot be positively identified. Some propose that proximity or adjacency implies connectivity. This is incorrect. We have previously found that proximity alone cannot predict connectivity, as neurons of the retina will traverse long stretches along a neighboring neuron without forming any sort of synapse, and not all densities are chemical or electrical synapses.[5] To precisely identify synaptic partners, our group uses multiple features present in ssTEM images of each synaptic type to accurately identify, annotate, and link pre- and post-synaptic pairs. We require evidence of synaptic vesicles, pre- and post-synaptic membranes to definitively identify synaptic contacts.

S.1 Conventional Synapses

Conventional synapses are the most common anatomical type of synapses found within the nervous system. As illustrated in figure S1A, the presynaptic side of the synapse is made up of a vesicle cloud made of numerous vesicles ranging in size between 30 and 50 nm. Often in TEM micrographs, the vesicle cloud is surrounded by an observable density (presumably from presynaptic protein machinery), indicated by a shadow in Figure S1A, and present in Figure S1B. Directly across the synaptic cleft (between 20-40nm) is the post synaptic density, which is darker and more uniform than the surrounding membrane. In order for a conventional synapse to be annotated in our volumes a vesicle cloud and post-synaptic density must be present for that synapse to be annotated and linked. The other features such as sizing and presynaptic densities often assist in the identification of these structures, however the vesicle cloud and post-synaptic density are necessary for final determination.

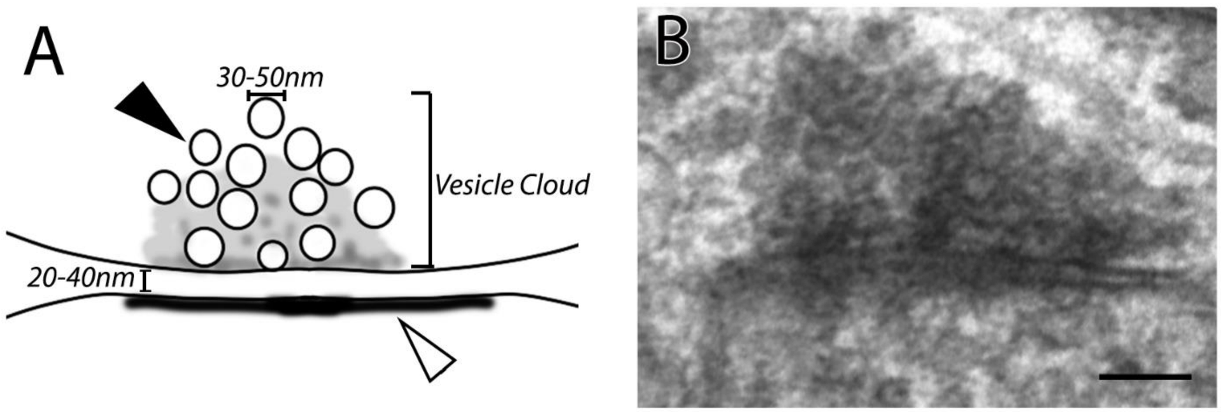


*Figure S1: Conventional Synapses (A) Schematic diagram of the primary features of a conventional synapse in a TEM micrograph. Arrows indicate vesicles (black) and the post-synaptic density (white). (B) TEM image of a conventional synapse in RPC1. Scale bar 100nm*

S.2 Ribbon Synapse

Ribbon synapses are specialized glutamatergic synapses found in the inner-ear hair cells and in the retina are found in photoreceptors and bipolar cells. As illustrated in figure S2A, the presynaptic side has a ribbon density (lamellae) with numerous tethered vesicles. The ribbon density can vary widely in length, and in RPC1 we find variations in shape, therefore we specifically look for densities with tethered vesicles to identify the ribbon synapse. Where the ribbon contacts the membrane, there is often a lighter density at its attachment point. Vesicles and the synaptic cleft are within the same size ranges as found in conventional synapses. The postsynaptic density in the IPL is often more prominent that that observed in conventional synapses, though it can be substantially smaller in the OPL opposing photoreceptors. It is also common to find a single ribbon presynaptic to multiple targets, this makes the presence of both a ribbon and the post-synaptic density(ies) critical for ribbon synapse and target(s) identification.

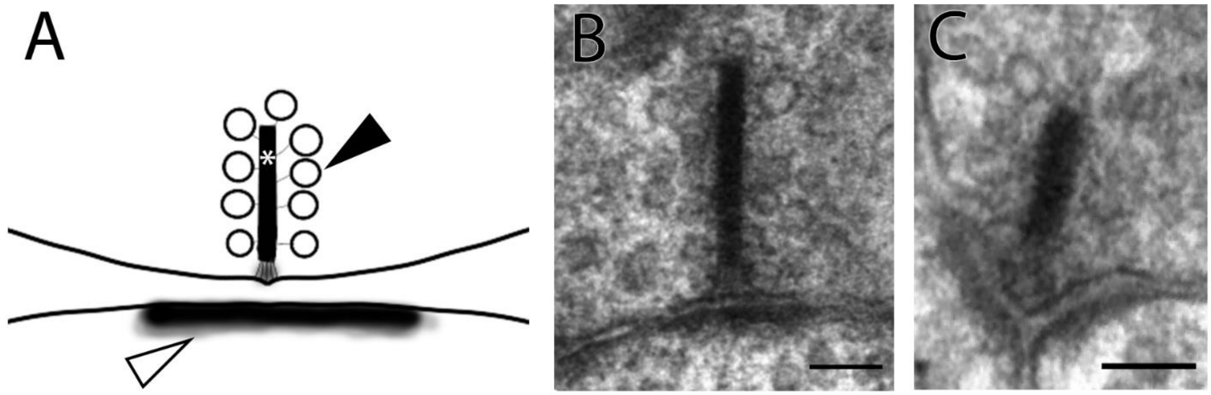


*Figure S2: Ribbon Synapses (A) Schematic diagram of the primary features of a ribbon synapse in a TEM micrograph. Arrows indicate vesicles (black) and the post-synaptic density (white). Astrisk highlights the ribbon density (B) TEM image of a ribbon*

*synapse in RPC1 opposing a single target. Scale bar 100nm (C) TEM image of a ribbon synapse in RPC1 opposing 2 target post-synaptic densities. Scale bar 100nm*

### S.3 Gap Junctions

Gap junctions are electrical and metabolic couplings between cells. They are found in many cell classes throughout the nervous system including retina and have multiple roles in visual processing, however they can vary dramatically in size and are often missed or misidentified when sufficient resolution is not present. As illustrated in figure S3, there is a density where 2 membrane densities come into direct opposition with one another with regular spacing between them, due to the conformation of connexins forming the gap junctions. In the middle of this space a third density is observable forming a pentalaminar conformation. The presence of the pentalaminar structure is the only full confirmation of gap junctions. There are also asymmetric, lighter, cytoplasmic densities found neighboring the more prominent. At our routine resolution (2.18nm/px), gap junctions appear as symmetrical regions of dark membrane that appear to "pinch" together, with small punctate supporting structures dotting the density. Gap junctions can be identified in our volumes, but if they are not perpendicular to the plane of section (Figure S3B) full confirmation may require recapture at higher magnification with goniometric tilt as was shown in figure 4.

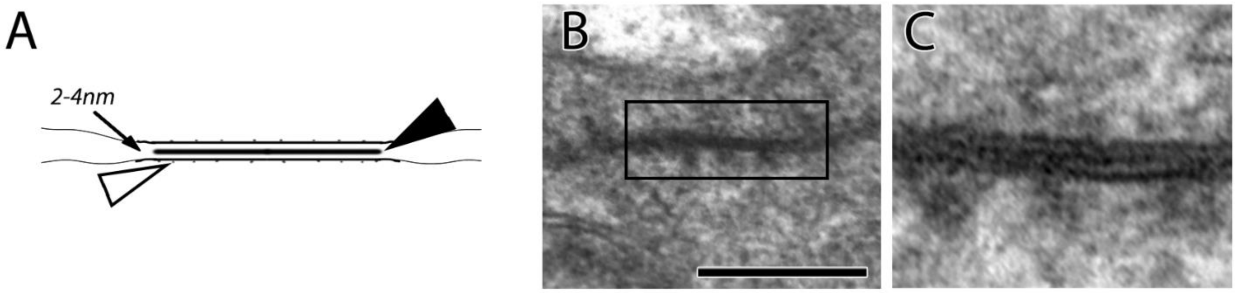


*Figure S3: Gap Junctions (A) Schematic illustration showing primary features of gap junctions in TEM. Arrowheads point to the center density unique to gap junctions (black) and the cytoplasmic densities associated with gap junctions (white). Arrow is pointing the intercellular space within the gap junction and its characteristic size. (B) TEM image of gap junction from RPC1 which is perpendicular to the plane of section allowing us to see the pentalaminar structure at 2.18nm/px. Scale bar 250nm. (C) Recaptured at 25k image of region outlined by the rectangle in B, contrast-adjusted to facilitate easier identification of the pentalaminar structure.*
